# Fix or Freeze? Spectral Differences Arising from Tissue Preparation in Chemical Imaging

**DOI:** 10.1101/2025.11.19.689284

**Authors:** Tianyi Zheng, Wihan Adi, Paul J Campagnola, Filiz Yesilkoy

## Abstract

Spectrochemical imaging has emerged as a powerful, label-free modality for visualizing the biochemical composition of tissues based on intrinsic vibrational signatures. Specifically, mid-infrared spectrochemical imaging (MIRSI) is becoming essential for fundamental biomedical research studying disease mechanisms, identifying biomarkers, and guiding drug development. However, the sensitivity of MIRSI to sample preparation protocols and its impact on spectral data interpretation remain poorly characterized. Here, we systematically compared spectral data collected from rat kidney and liver tissues prepared using standard fresh frozen (FF) and formalin-fixed, paraffin-embedded (FFPE) tissue processing methods using quantum cascade laser (QCL)–based MIRSI. We applied frequently used spectral data processing techniques, including uniform manifold approximation and projection (UMAP), correlation matrices, and second-derivative spectral analysis, to characterize preparation-induced differences. FF samples preserved a broader range of biochemical signals, retaining the innate chemical composition of tissues, while FFPE tissues showed reduced spectral diversity and absorption signal intensity. Moreover, a unique spectral band at 1026 cm^−1^, likely arising from formalin-induced collagen crosslinking, was identified specifically in FFPE samples. Our findings demonstrate that tissue preparation substantially alters chemical and morphological information captured by MIRSI, necessitating careful consideration of processing protocols in workflows involving chemical imaging and Artificial Intelligence (AI) - based spectral analysis.

## Introduction

Chemical imaging techniques are emerging as powerful analytical methods for tissue analysis in biomedical sciences. Specifically, mid-infrared spectrochemical imaging (MIRSI) is becoming essential for fundamental biomedical research studying disease mechanisms, identifying biomarkers, and guiding drug development ^1–3^. Recent advances in mid-infrared (mid-IR) photonics have improved the standard MIRSI instrumentation, such as Fourier transform IR (FTIR) spectrometers^4^, while introducing new spectrochemical imaging modalities, including dual-frequency combs^5^, quantum cascade laser (QCL)-based microscopes^6,7^ and optical photothermal IR (O-PTIR) imaging^8^. These MIRSI modalities provide spatially resolved molecular information in *ex vivo* tissue and cell investigations, revealing biological insights beyond the capabilities of optical imaging methods requiring contrast-generating agents. Unlike traditional histological methods that reveal information on structural and morphological features of tissues, chemical imaging captures molecular fingerprints imprinted in optical spectra that reveal the underlying biochemical differences in healthy and diseased tissues^9–11^.

The extent of information these techniques can provide in biomedical research and clinical settings depends on the preservation of intrinsic chemical signals. The mid-IR molecular fingerprint spectrum includes key vibrational bands, such as the amide I and II bands (∼1650 and ∼1550 cm^-1^) that carry information on proteins’ secondary structures, the lipid-associated bands (∼1700-1740 cm^-1^), and phosphoric acid or carbohydrate modes below 1400 cm^-1^, which can be used as biomarkers of functional and metabolic cell states. However, these spectral features are potentially sensitive to sample preparation protocols and can be masked or lost if the biochemical integrity of the tissue is compromised.

Two widely used preparation methods,fresh frozen (FF) and formalin-fixed, paraffin-embedded (FFPE), offer complementary strengths for different types of applications. FFPE ensures long-term preservation and maintains tissue morphology but induces chemical alterations through protein crosslinking and solvent-based dehydration^12-14^. In contrast, FF tissue preservation protocols better retain labile biomolecules but present logistical challenges related to storage and handling^15^. Previously, FTIR spectra from FF tissues were compared to those from FFPE and deparaffinized tissues, revealing spectral changes associated with the tissue preparation processes^16-18^. Similarly, comparative studies using Raman spectroscopy of dewaxed-FFPE and FF tissues from both cancerous and healthy samples reported decreased spectral signal intensity in FFPE samples^19^. However, these investigations were limited to ensemble-averaged FTIR or Raman spectra collected from large tissue areas rather than spatially resolved large datasets collected by spectrochemical imaging. Consequently, the effects of tissue preparation protocols on spectral datasets were observed but undermined, hindering the reliable interpretability and reproducibility of chemical imaging approaches across different biomedical research and clinical applications.

In this study, we investigate how sample preparation methods affect both the spectral and structural aspects of chemical tissue imaging using a QCL-based MIRSI microscope. By analyzing thousands of spectra from rat kidney and liver tissues prepared using standard FF and FFPE protocols, we show that FFPE processing causes significant spectral degradation, including attenuation or disappearance of spectral bands associated with functional biomolecules, such as lipids, nucleic acids, and proteins. Moreover, our second-derivative spectral analysis revealed decreased spectral band diversity in FFPE tissues. Importantly, we also identified an absorption peak at 1026 cm^−1^, which can be assigned to –CH_2_OH groups, uniquely in FFPE tissues. This peak likely arises from formaldehyde-induced crosslinking of collagen. To assess whether MIRSI spectra could be used to distinguish tissue types and preparation methods, we applied UMAP for unsupervised clustering, revealing that FF tissue spectra maintained biochemical diversity sufficient for tissue classification, while spectra from FFPE tissues formed a compressed cluster. Logistic regression on UMAP-projected data further revealed that discriminative features between FF and FFPE samples primarily originate from the glycogen and Amide regions. These results emphasize the critical role of tissue preparation in preserving the innate biochemical composition of tissues, highlighting the potential limiting effects of chemical fixing in MIRSI data interpretation using machine learning techniques in chemical imaging pipelines.

## Materials and methods

### Animal Handling and Tissue Collection

All procedures involving live rats were conducted in accordance with the NIH Guide for the Care and Use of Laboratory Animals and were approved by the Institutional Animal Care and Use Committee (IACUC) at the University of Wisconsin School of Medicine and Public Health. Adult rats were euthanized via intraperitoneal injection of a mixture containing ketamine (100 mg·kg^−1^), xylazine (20 mg·kg^−1^), and acepromazine (3 mg·kg^−1^), followed by transcardial perfusion with phosphate-buffered saline (PBS) and 4% paraformaldehyde (PFA). Liver and kidney tissues were immediately dissected and divided in half for parallel post-processing by either fresh freezing in OCT or FFPE protocols.

### Tissue Embedding and Sectioning

For FF tissue preparation, liver and kidney tissues were transferred to cryomolds, embedded in OCT compound, and snap-frozen on dry ice. Frozen tissues were sectioned at 10 µm thickness using a sliding microtome and immersed briefly in deionized water to dissolve residual OCT compound. For FFPE processing, the other half of the kidney and liver tissue samples were post-fixed overnight in 4% PFA, dehydrated, and embedded in paraffin wax. Paraffin-embedded samples were also sectioned at 10 µm using the same microtome and subsequently deparaffinized using standard xylene and ethanol washes. All tissue sections were mounted onto pristine calcium fluoride (CaF_2_) substrates and stored at 4 °C until spectral imaging within 24h.

### Quantum Cascade Laser–Based Mid-Infrared Imaging

Mid-infrared hyperspectral imaging was performed using a Spero-QT microscope (Daylight Solutions, Inc.) equipped with four QCL modules, providing spectral coverage from 950 to 1800 cm^−1^ (Figure 1 b), as described in our previous work^20^. High-magnification scans were acquired using a 12.5× infrared objective (NA = 0.7; pixel size = 1.3 µm) and an uncooled microbolometer focal plane array (480 × 480 pixels). All spectral data were collected at a spectral resolution of 2 cm^−1^.

**Figure 1.**
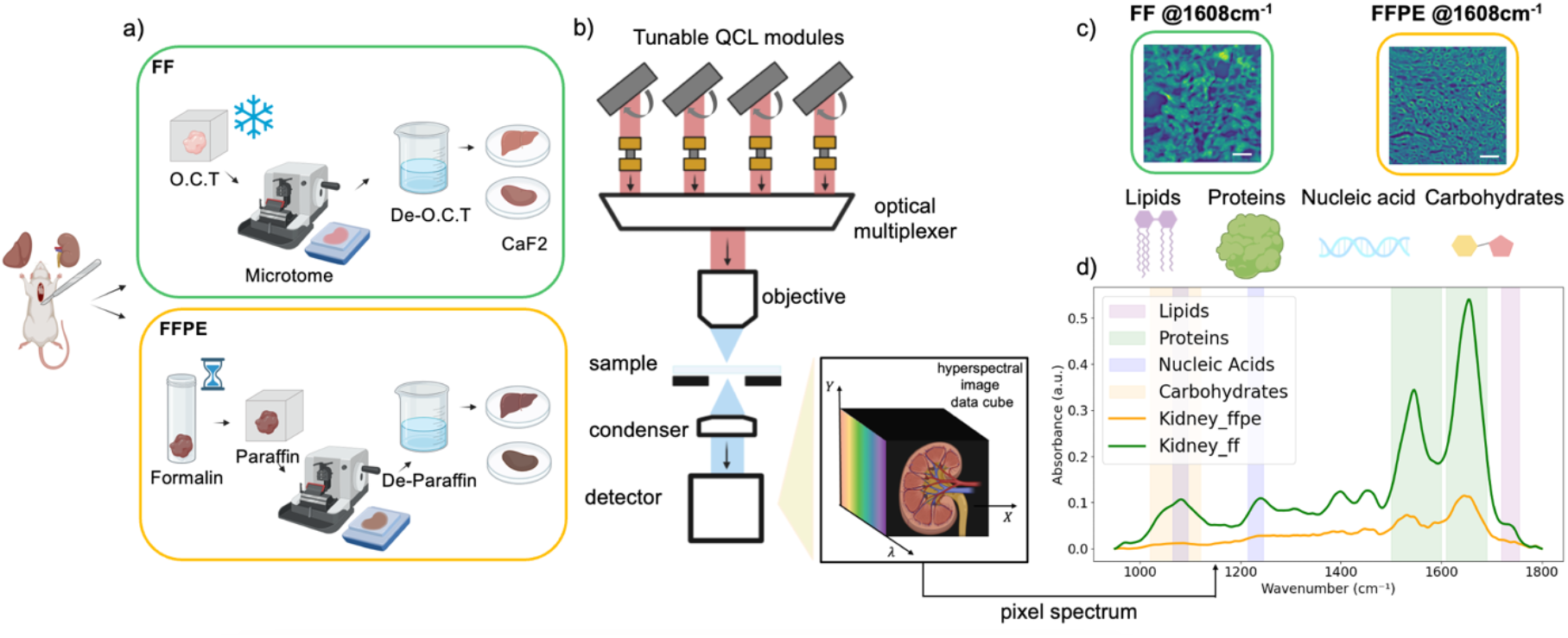
Sample preparation and mid-infrared spectrochemical imaging workflow. a). Schematic of tissue processing for MIRSI. Rat kidney and liver tissues were prepared either by fresh freezing (FF) in optimal cutting temperature (OCT) compound or formalin fixation followed by paraffin embedding (FFPE). All tissues were sectioned at 10 µm thickness and mounted on CaF_2_ substrates. Before imaging, the embedding media, OCT or paraffin, were removed. b). Schematic of the QCL-based MIRSI system used for hyperspectral imaging in widefield transmission mode across fingerprint spectral range of 950–1800 cm^−1^. c). Representative mid-IR images of FF and FFPE kidney tissues captured at the Amide I region (1608 cm^-1^). Scale bars indicate 100 μm. d). Measured absorbance spectra from FF and FFPE kidney tissues. Each spectrum was averaged from ∼100,000 pixels. The color bands in the background highlight spectral regions associated with different biomolecules.

### Preprocessing of Spectrometric Data

After hyperspectral 3D data collection, datasets were processed using custom Python scripts (Figure 2). For each sample type (FF or FFPE liver or kidney; four types in total), ten regions of interest (ROIs) of 0.650 × 0.650 mm^2^ (480 × 480 spectra per ROI) were acquired. A total of eight ROIs per sample type were retained for further analysis, yielding 8 × 480 × 480 ∼ 1.8 M spectra in total per sample.

**Figure 2.**
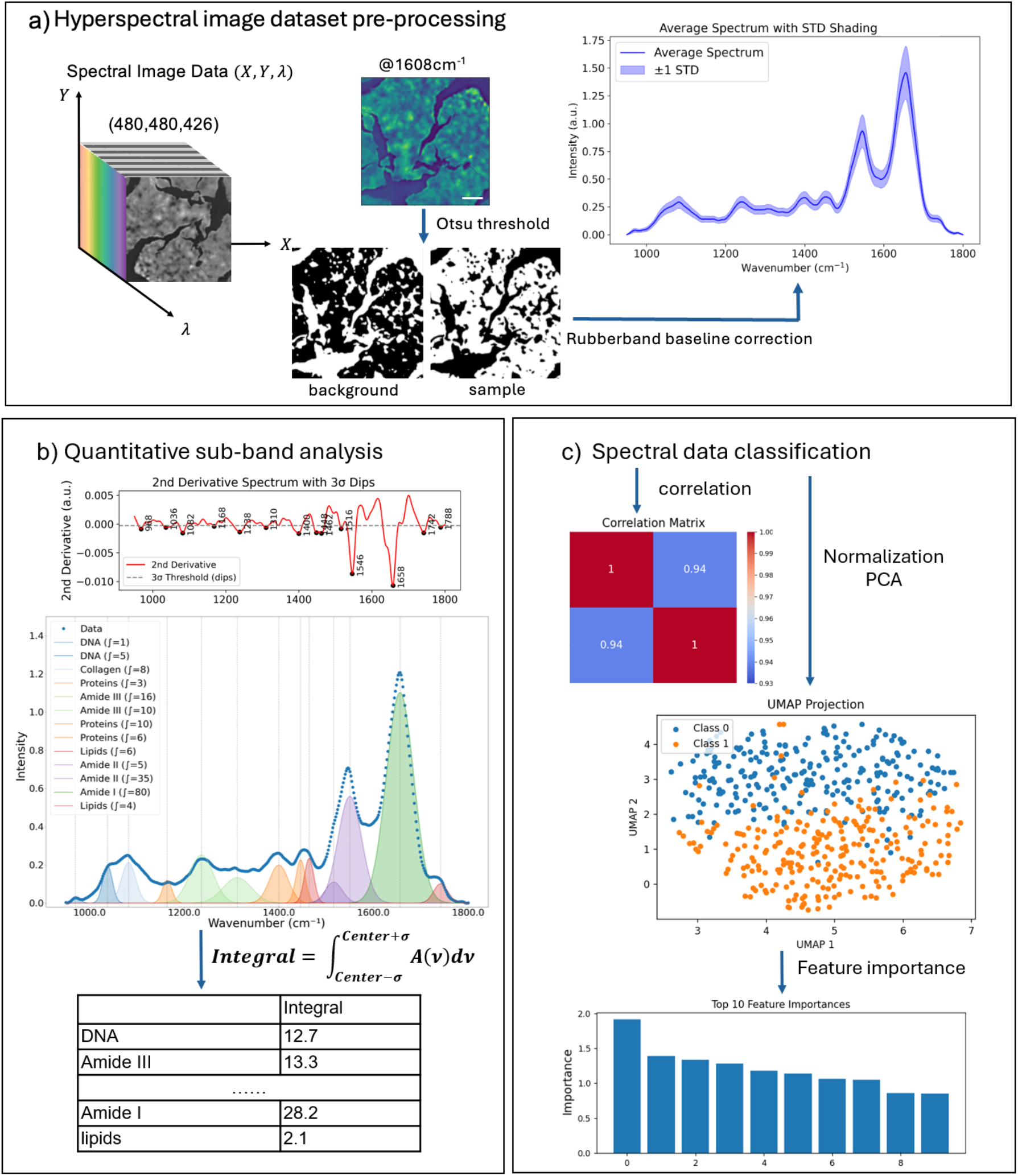
Overview of spectral data processing methods. a). Eight ROIs were selected from each tissue sample. Tissue-associated sample spectra were extracted by applying Otsu thresholds to a single mid-IR image collected at 1608 cm^-1^, Amide I region. Scale bars indicate 100 μm. Next, rubberband baseline correction was applied to the extracted spectra to remove scattering contributions. b). For quantitative sub-band analysis, second-order Savitzky– Golay derivatives were applied to the filtered, polymer-fitted spectral signal. Functional vibrational bands were identified using 3σ rule and quantified using composite Simpson’s integration. c). For spectral data classification, correlation heatmaps were generated to illustrate similarities between and within samples. Absorption spectra were Z-score normalized and then subjected to PCA for dimensionality reduction. The processed spectra were classified by unsupervised ML methods (UMAP), which was followed by logistic regression to evaluate spectral feature importance.

Before spectral data preprocessing, we first identified blank pixels, where there is no tissue present, and removed them from the datasets of interest. Those off-tissue pixels in each ROI were identified using Otsu thresholding^21^ applied to the mid-IR images collected at the 1608 cm^−1^, Amide I band, primarily the strongest band in the spectrum. The average of the discarded off-tissue pixels was set as the background spectrum to calculate the noise. Next, we took the second derivative of the background spectrum and calculated its standard deviation (σ), which was then set as the background noise and used to distinguish significant absorption peaks from the background.

From each ROI, 2,000 tissue-associated pixels were randomly selected and saved for further analyses, resulting in a dataset of 16,000 spectra per sample type. Each tissue-associated pixel spectrum was preprocessed by applying a rubberband baseline correction to remove the scattering contributions^22^.

### Quantitative sub-band analysis of spectrometric data

To enhance absorbance band resolution and minimize baseline variations, the second derivative of each absorbance spectrum was computed using a Savitzky–Golay filter^23^ (window length = 13, polynomial order = 2; scipy.signal) (Figure 2b). Due to edge effects from filtering, 26 spectral points were trimmed from the original 426-point spectral range. This procedure sharpened overlapping vibrational bands, abundant in complex biological samples, and improved peak detection. From the second-derivative spectra, significant peaks were identified using the 3σ rule, which identifies significant peaks and sub-peaks lying outside the calculated 3σ range^24^. Here, σ was calculated from the discarded off-tissue background spectra, ensuring that peak detection was grounded in statistical rigor rather than an arbitrary threshold.

From each ROI, a subset of 100 pixel-spectra was randomly selected for peak occurrence analysis, and the average spectra were used for sub-band Gaussian fitting (Figure 2b). To quantify the occurrence frequency of specific vibrational bands, peak occurrence was calculated as the number of pixels within each ROI that contains a significant peak (a band passing the 3σ rule) at a spectral position with ± 2 cm^−1^ tolerance. The total occurrence of each vibrational band was then expressed as a count across all analyzed spectra across the ROIs, and the results were plotted as box-and-whisker plots in Figures 3 and 4. This approach reflects how often a characteristic absorbance feature was present among different samples, providing a statistical representation of spectral consistency and biochemical prevalence within the tissue regions.

**Figure 3.**
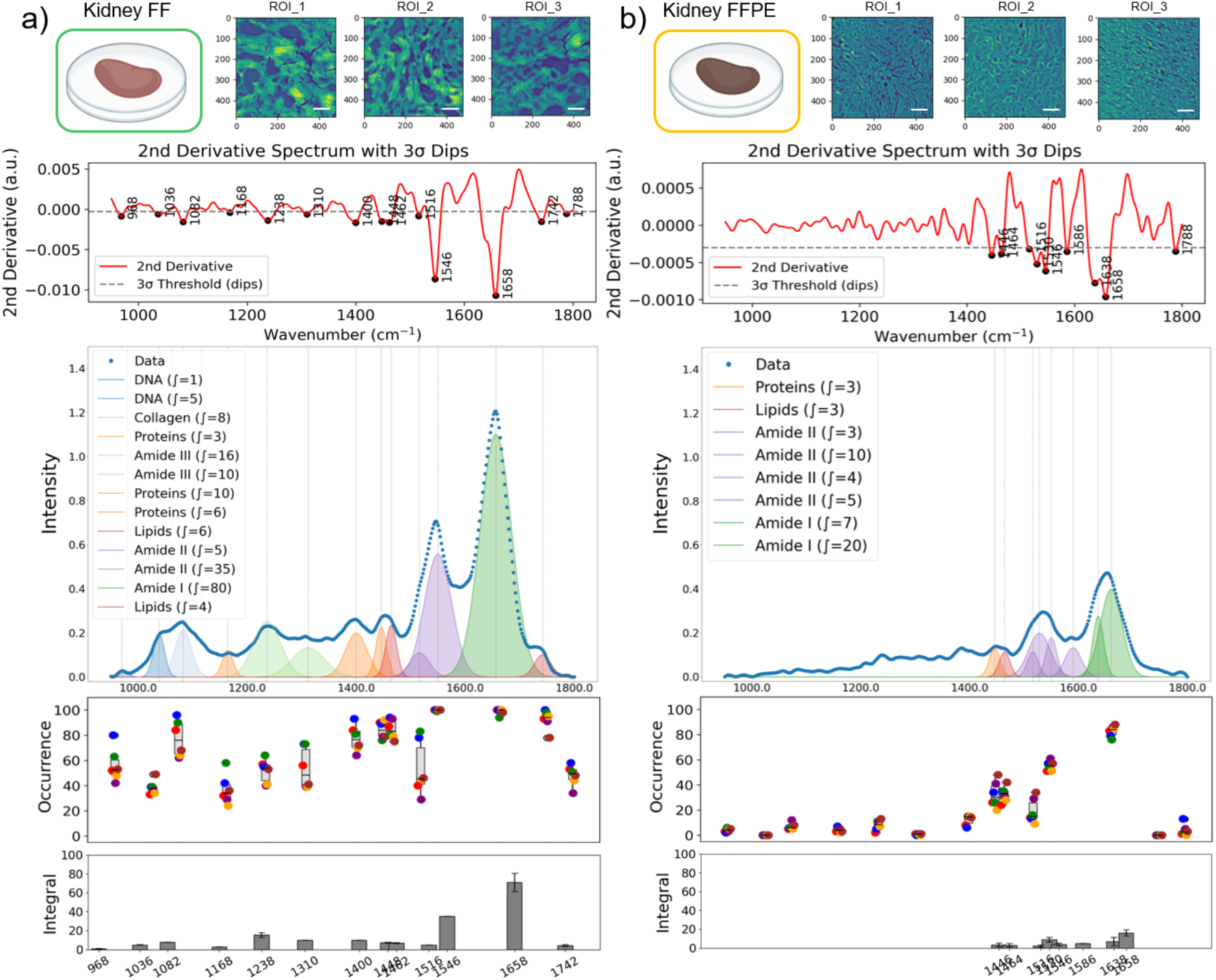
Comparative spectral data variations in kidney FF and FFPE tissues. Three representative mid-infrared images (ROIs) captured at the Amide I band (1608 cm^-1^) are shown for kidney FF (a) and FFPE (b) tissues. Scale bars indicate 100 μm. Second-derivative spectra with dips detected by the 3-sigma rule. The intensity plots show the average mid-infrared spectra generated by randomly selected 120 pixel-spectra from eight ROIs, and the sub-bands were identified by second-derivative analysis, and Gaussian fitting was applied for FF (a) and FFPE (b) tissues. The occurrence of the detected absorbance peaks from eight ROIs was plotted (each ROI was represented by 100 pixels) for FF (a) and FFPE (b) tissues. Each data point represents the average occurrence from a single ROI; the grey box-and-whisker plot shows the center and spread of the peak occurrence. In the bottom integral plots, the dark grey bars represent the average and standard deviation of the integral calculated for each specific band.

**Figure 4.**
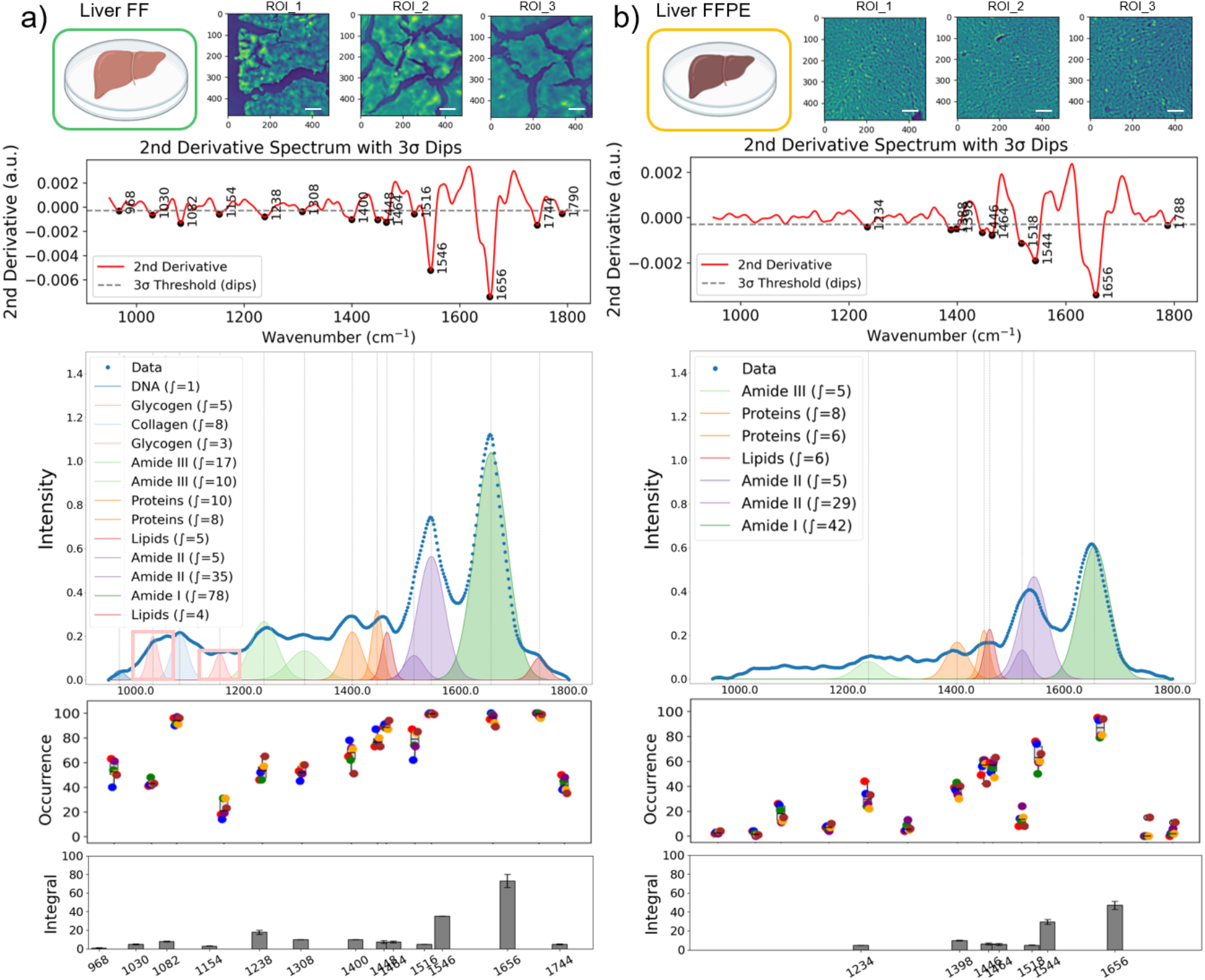
Comparative spectral data variations in liver FF and FFPE tissues. Three representative mid-infrared images captured at the Amide I band (1608 cm^-1^) are shown for liver FF (a) and FFPE (b) tissues. Scale bars indicate 100 μm. Second-derivative spectra with dips detected by the 3-sigma rule. The intensity plots show average mid-infrared spectra generated by randomly selected 120 pixel-spectra from eight ROIs, and the sub-bands were identified by second-derivative analysis, and Gaussian fitting was applied for FF (a) and FFPE (b) tissues. The occurrence of the detected absorbance peaks from eight ROIs was plotted (each ROI was represented by 100 pixels) for FF (a) and FFPE (b) tissues. Each data point represents the average occurrence from a single ROI; the grey box-and-whisker plot shows the center and spread of the data. In the bottom integral plots, the dark grey bars represent the average and standard deviation of the integral calculated for each specific band.

Quantitative absorbance information for each sub-band was extracted by numerical integration within defined spectral windows using the composite Simpson’s rule (scipy.integrate)^20^, and vibrational bands were assigned based on established spectral positions of functional groups^25^. The spectral regions used for integration are listed in Table 1.

**Table 1.** Reference integration intervals along second-order derivative absorption spectra.

| Assignment | Peak center [$\text{cm}^{-1}$] | Reference |
| --- | --- | --- |
| DNA | ~966 | 26 |
| Glycogen | ~1030 | 27,28 |
| DNA | ~1035 | 29 |
| DNA | ~1082 | 30 |
| Glycogen | ~1154 | 31 |
| Proteins | ~1168 | 32 |
| Amide III | ~1238 | 33 |
| Amide III | ~1308 | 34 |
| Proteins | ~1400 | 35 |
| Proteins | ~1448 | 27 |
| Lipids | ~1462 | 26 |
| Amide II | ~1516 | 36 |
| Amide II | ~1546 | 36 |
| Amide I | ~1658 | 37 |
| Lipids | ~1744 | 34 |

### Spectral data classification

For spectral data classification, a random subset of 2,000 pixel-spectra per ROI (160,000 spectra in total across sample types) was selected. Correlation between ROIs was calculated and displayed in a heatmap, illustrating the degree of similarity within ROIs of the same sample type and highlighting differences across distinct samples. Absorbance spectra were Z-score normalized (mean = 0, standard deviation = 1) for standardization, then subjected to principal component analysis (PCA) for dimensionality reduction, followed by UMAP for visualization of group separation. Finally, feature importance was evaluated using logistic regression, and the top 40 informative wavenumbers were identified (Figure 2c).

## Results and discussion

### Effect of Sample Preparation on Spectral Integrity in Kidney and Liver Tissues

To evaluate the impact of tissue sample preparation on mid-infrared spectral datasets, we analyzed rat kidney and liver tissues processed using either FF or FFPE protocols separately (Figures 3, 4). All measured tissues were sectioned at 10 µm thickness using the same microtome within 24 hours to minimize light-path-dependent signal variations in the absorbance spectra. In kidney tissues, FF sections consistently exhibited stronger overall absorption and greater spectral diversity (higher occurrence) compared to FFPE tissues (Figure 3). To support our empirical observations with rigorous data analysis, we considered spectral data from eight ROIs (100 spectra per ROI) per FF and FFPE samples. Based on our 3σ rule, thirteen distinct peaks were consistently observed in FF kidney tissues, with robust preservation of DNA, protein, and lipid-associated bands (Figure 3a). In contrast, only eight bands were detected in FFPE tissues (Figure 3b). Although protein-associated bands (Amide I and II) remained detectable, signals from DNA and lipids were significantly reduced or absent. The overall diversity of spectral bands was summarized in the occurrence plots in Figure 3a and b for FF and FFPE samples, respectively. Variations in the band occurrence across spectra from eight different ROIs can be attributed to spatial compositional changes across the sectioning cross-section of the kidney, consisting of different tissue and cell types. Moreover, on average, the absorbance of the Amide 1 band in FFPE was 66.3% lower ((*Integral*_*FF*_−*Integral*_*FFPE*_)/*Integral*_*FF*_) than the FF kidney tissue measurements based on our band integral analysis.

We hypothesize that the FFPE protocol is chemically harsh: repeated xylene and ethanol washes can extract or degrade biomolecular components, leading to lower signal intensity and fewer detectable peaks. Similar observations have been reported previously^16,17,19^ using various chemical imaging techniques. Notably, our spectral imaging approach captures structural and spatially resolved spectral changes due to tissue post-processing protocols on the key compositional information captured by chemical imaging methods.

Similarly, FF liver tissues showed more diverse and stronger molecular signatures across the spectral range compared to FFPE tissue sections (Figure 4). While Amide I and II bands, which are typically strong, remained detectable in FFPE liver tissues, multiple bands below 1400 cm^−1^ were lost. Considering spectral data from eight ROIs (100 spectra per ROI) per FF and FFPE samples, we consistently identified thirteen distinct peaks in FF liver tissues, while only seven peaks were captured across different pixel spectra in FFPE samples. The occurrence of significant peaks was presented for FF and FFPE in Figure 4a and b, respectively, summarizing the diversity of bands captured from different ROIs. The variations of the peak occurrence across the ROIs can be attributed to spatial compositional variations in the liver, typically hosting a number of different tissue layers and cell types. We also identified that the average absorbance of the Amide 1 band in FFPE was 46.2% lower than the FF liver tissue measurements, in alignment with our findings with kidney tissue.

The liver is an organ with a high metabolic complexity and an abundance of chemically labile molecules. The reduction in spectral intensity and band loss in FFPE samples can be attributed to the chemical consequences of formalin fixation, including protein crosslinking as well as removal of lipophilic and labile components during dehydration in ethanol and paraffin embedding. For example, the glycogen band (@1030cm^-1^), a marker of hepatic metabolism^38^, was only observed in FF liver tissues (as highlighted in Figure 4a). Collectively, these findings underscore the limitations of FFPE for comprehensive chemical mapping and highlight the advantages of FF preparation in preserving the biochemical complexity of metabolically active tissues for MIRSI analysis.

### U-map and Feature Importance Analysis of Spectral Profiles

To further investigate the impact of sample preparation on the discriminatory power of spectral features, we first calculated correlation matrices between ROIs from different sample types and displayed them as a heatmap (Figure 5a). The correlation matrices illustrate a high degree of similarity in spectra from different ROIs of the same sample type, while highlighting differences due to tissue processing for both liver and kidney tissues.

**Figure 5.**
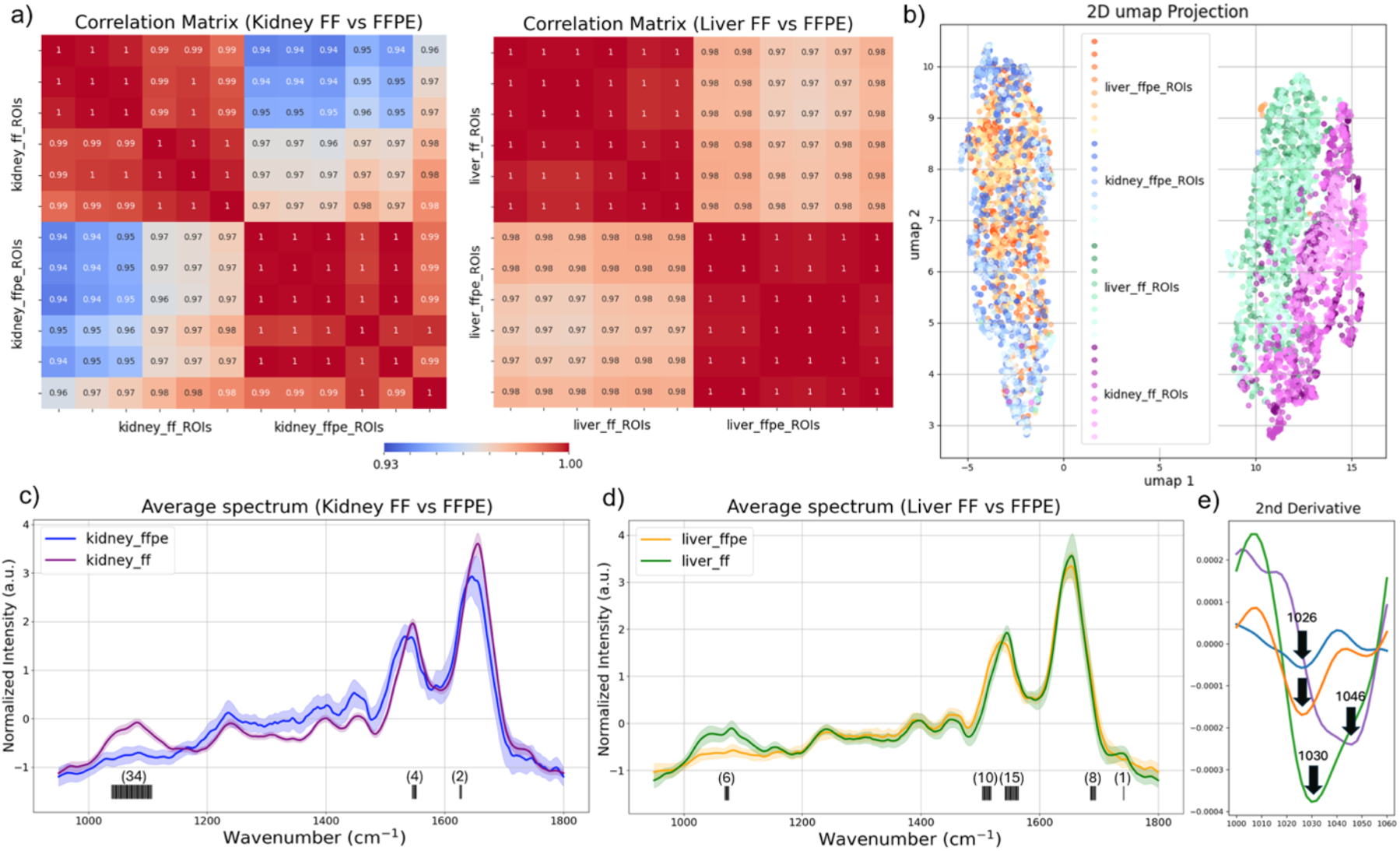
Correlation matrices and UMAP clustering reveal tissue processing effects on the spectral discrimination power in chemical imaging. a). Correlation heatmaps demonstrate the degree of spectral similarity within ROIs of the same sample type and highlight differences across tissues prepared using different protocols. b) UMAP projection of normalized spectra from FF and FFPE rat kidney and liver tissues. c) Averaged and normalized spectra from liver FF and FFPE samples were plotted and top 40 discriminative wavenumbers were identified and indicated as the black lines under the spectra. d) Averaged and normalized spectra from kidney FF and FFPE samples were plotted and top 40 discriminative wavenumbers were identified and indicated as the black lines under the spectra. Number in the parentheses indicate the number of important features detected. e) Zoom-in second derivative spectra from 1000 cm^-1^ to 1060 cm^-1^ showing the distinct absorption band at 1026 cm^-1^ detected only in FFPE samples.

Moreover, we applied UMAP to preprocessed spectra from rat kidney and liver tissues. A total of 20,000 spectra were used for UMAP after reducing spectral dimensionality using PCA. From a total of eight ROIs, 5000 spectra were randomly selected from each sample type (FF and FFPE liver and kidney). UMAP revealed distinct clustering patterns based on the tissue processing method. Independent of tissue type, liver or kidney, spectra from FFPE-processed samples formed a single, tightly grouped cluster. On the contrary, spectra from FF samples retained distinct spectral features between kidney and liver tissues (Figure 5b). Given that the considered organs, the liver and kidney, have distinct biological and physiological functionalities, the discrimination between their chemical composition captured by the FF tissue spectra is expected. On the contrary, the overlapping UMAP clusters among the FFPE tissue spectra indicate that the chemically intrusive sample processing removes the key spectral features unique to each organ type.

To identify the most discriminative spectral features, we applied a logistic regression model to the UMAP-clustered spectral datasets and identified the highest-ranked feature importance wave-numbers across kidney and liver samples. The top 40 contributing wavenumbers were predominantly located in the lower wavenumber region (nucleic acids bands, carbohydrate-associated bands), as well as the Amide I and II regions, which are the key vibrational modes linked to functional biomolecules (Figure 5c, d). These results highlight the superior spectral fidelity of FF tissues in preserving biologically relevant features, which are essential to apply chemical imaging for biomedical research and diagnostic applications.

### Additional Spectral Feature in FFPE Tissues

In addition to signal loss, FFPE processing introduced a distinct absorption band at 1026 cm^−1^, which was absent in all FF samples (Figure 5e). This peak, consistently detected in FFPE liver sections and partially in kidney, can be assigned to –CH_2_OH stretching vibrations, likely arising from formaldehyde-induced hydroxymethylation of collagen and other hydroxyl-containing residues. Prior studies have shown that formalin can modify amino acids such as lysine and hydroxyproline, forming hydroxymethyl derivatives and methylene bridges^12^.

The presence of an additional peak suggests that FFPE not only diminishes native molecular features but also introduces fixation-specific artifacts. Moreover, Figure 5c and d indicates a spectral shift in Amide I and II bands, which can be attributed to protein crosslinking and structure modifications of the fixing process. In a hypothesis-driven biomedical study, such chemically altered signatures may interfere with spectral interpretations. Our work highlights the need to account for tissue preparation-induced changes in chemical imaging workflows.

### Implications for Chemical Imaging-Based Tissue Analysis

Together, our findings highlight the important role of sample preparation protocols on the fidelity and interpretability of spectrochemical tissue image datasets. While FFPE is widely used for its morphological preservation and long-term storage benefits, the harsh chemical treatments involved can compromise labile molecular signatures and introduce fixation-specific artifacts, limiting analytical depth and complicating AI-based classification or biomarker discovery pipelines. In contrast, FF preparation, though more logistically demanding, preserves both biologically and physiologically informative molecular signatures. However, as the mid-IR images show, the tissue morphology is hindered in FF sections. This is not a significant hurdle because morphology can be easily analyzed using well-established histopathology techniques using accessible optical widefield microscopes. As MIRSI applications continue to expand in biomedical research, careful consideration of sample preparation protocols will be essential to maintain the innate chemical composition, which can enable accurate spectral classification and support robust machine learning applications.

## Conclusion

This study demonstrates that tissue sample processing significantly affects the preservation of biochemical information in mid-IR spectrochemical imaging. By directly comparing mid-IR spectra from FF and FFPE rat kidney and liver tissues, we show that FF samples consistently preserved a more diverse and intense range of biochemical signals, compared to that of FFPE tissues. Notably, an absorption feature at 1026 cm^−1^, which can be attributed to –CH_2_OH groups formed via formaldehyde-induced collagen crosslinking, was detected exclusively in FFPE tissues. These findings highlight that FFPE processing not only diminishes native molecular signatures but can also introduce fixation-specific artifacts. As chemical imaging advances toward clinical and AI-integrated biomedical applications, optimizing and standardizing sample preparation across the chemical imaging research community will be essential to ensure spectral fidelity and accurate biochemical interpretation.

## Acknowledgement

The authors thank the University of Wisconsin Translational Research Initiatives in Pathology (TRIP) Laboratory, supported by the UW Department of Pathology and Laboratory Medicine, UWCCC (P30 CA014520), and the Office of the Director-NIH (S10 OD023526) for use of its facilities and histology services. F.Y. and P.C. acknowledge financial support from the U.S. National Institutes of Health (R61CA281795).

## Notes

### Competing Interest Statement

The authors have declared no competing interest.

